# A novel automated approach for improving standardization of the marble burying test enables quantification of burying bouts and activity characteristics

**DOI:** 10.1101/2021.10.16.464233

**Authors:** Lucas Wahl, A. Mattijs Punt, Tara Arbab, Ingo Willuhn, Ype Elgersma, Aleksandra Badura

## Abstract

The marble burying test is a commonly used paradigm to screen phenotypes in mouse models of neurodevelopmental and psychiatric disorders. The current methodological approach relies solely on reporting the number of buried marbles at the end of the test. By measuring the proxy of the behavior (buried marbles), rather than the behavior itself (burying bouts), many important characteristics regarding the temporal aspect of this assay are lost. Here we introduce a novel, automated method to quantify mouse behavior throughout the duration of the marble burying test with the focus on the burying bouts. Using open-source software packages, we trained a supervised machine learning algorithm (the “classifier”) to distinguish burying behavior in freely moving mice. In order to confirm the classifier’s accuracy and uncover the behavioral meaning of the marble burying test, we performed marble burying test in three mouse models: *Ube3a^m-/p+^* (Angelman Syndrome model), *Shank2*^-/-^ (autism model), and *Sapap3*^-/-^ (obsessive-compulsive disorder model) mice. The classifier scored burying behavior accurately and consistent with the literature in the *Ube3a^m-/p+^* mice, which showed decreased levels of burying compared to controls. *Shank2*^-/-^ mice showed a similar pattern of decreased burying behavior, which was not found in *Sapap3*^-/-^ mice. Tracking mouse behavior throughout the test enabled us to quantify activity characteristics, revealing hypoactivity in *Ube3a^m-/p+^* and hyperactivity in the *Shank2*^-/-^ mice, indicating that mouse activity is unrelated to burying behavior. Together, we demonstrate that our classifier is an accurate method for the analysis of the marble burying test, providing more information than the currently used methods.

**Significance Statement:** The marble burying test is widely used in phenotyping neurodevelopmental and neuropsychiatric disorder mouse models. Currently, its analysis consists of manually scoring the number of buried marbles upon the completion of the assay. This approach is not standardized across laboratories, and leaves out important variables such as movement characteristics and information about the burying bouts. This leads to divergent interpretations of the marble burying test, ranging from anxiety to cognitive impairment. We introduce a method that reliably tracks mouse behavior throughout the experiment, classifies the duration and number of the burying bouts, and is generalizable across laboratories. Using machine learning for measuring the actual burying behavior standardizes this method, and provides rich information about the burying characteristics.

## Introduction

The marble burying test (Pinel & Treit, 1978) is a commonly used paradigm, aimed at studying repetitive behavior as well as an anxiety-like phenotype (Angoa-Pérez et al., 2013; Broekkamp et al., 1986; Thomas et al., 2009). It has more recently been used to study models of neuropsychiatric and neurodevelopmental disorders, with 87.82% of studies being done in mice (Çalişkan et al., 2017). The behavioral meaning of marble burying is however highly debated throughout literature. Studies have found that burying behavior can be selectively inhibited by anxiolytics and antidepressants (Ichimaru et al., 1995; Nicolas et al., 2006). However, results show no correlation with “anxiety-related” responses in the open-field or light–dark tests (such as elevated plus maze), nor are they correlated with overall exploratory activity (Thomas et al., 2009). Furthermore, marble burying and digging-event frequency were found to correlate only on the first 2 out of 5 days of repeated testing, showing a dissociation between burying and digging behavior (Thomas et al., 2009).

Our understanding of the behavioral relevance of the marble burying test is hampered by the conventional scoring methods that fail to analyze burying behavior as a whole, and rather focus only on the number of buried marbles. In the classical approach, mice are removed from the apparatus at the end of testing and an experimenter assesses visually how many marbles are covered more than a chosen threshold. This threshold varies throughout literature but is commonly set at a marble being either 50% or two-thirds covered by bedding to be considered “buried” (Angoa-Pérez et al., 2013; Kalariya et al., 2015; Sonzogni et al., 2019; Thomas et al., 2009). The main benefit of this method is high throughput due to the short analysis time needed. It has also been shown to be consistent within a given mouse model (Sonzogni et al., 2018). However, most laboratories only use visual scoring of buried marbles as a proxy for burying behavior, which fails to include many of the features of actual behavior in the analysis, disregarding possible changes in burying patterns that are not visible in the end result. Rare studies make use of images or video recordings, though analysis is often still done with manual annotation of frames (Serra et al., 2021). This method is highly time-consuming and is therefore not widely adopted.

Automated classification of behaviors based on machine learning algorithms provides a way to study actual behaviors over time, instead of a single parameter end result (Kabra et al., 2013; Pereira et al., 2019; Wiltschko et al., 2020). In the last few years, this technique has been used to identify social and locomotor behaviors of fruit flies (Aso et al., 2014), grooming events in mice (Boom et al., 2017) and encounter behaviors in honeybees (Blut et al., 2017), making the analysis easily transferable across laboratories. Although lower throughput than visual inspection, the collected videos can be analyzed in a batch mode, significantly speeding up the analysis process.

Additional information regarding the ways in which mice alter their burying characteristics might elucidate the behavioral meaning of marble burying behavior in a way that is impossible to find in only scoring the resulting buried marbles. Here, we used supervised machine learning to train a classifier in order to provide a method for repeatable inter- and intra-experimenter scoring of burying behavior that gives additional information regarding spatial and temporal burying characteristics. To test the accuracy of the classifier we used a mouse model for Angelman Syndrome (*Ube3a^m-/p+^*), which has an established burying phenotype, to validate our scoring results with literature (Huang et al., 2013; Sonzogni et al., 2018; Wang et al., 2018).

Furthermore, we aimed to elucidate the behavioral meaning of marble burying by utilizing two additional selective mouse models known for repetitive and compulsive/anxious behaviors: the *Shank2*^-/-^ model of autism spectrum disorder (ASD; Schmeisser et al., 2012; Won et al., 2012) and the *Sapap3*^-/-^ model of obsessive-compulsive disorder (OCD; Welch et al., 2007), respectively. Using these models, we aimed to determine whether marble burying can be considered a purely repetitive or a compulsive behavior. We show that unlike grooming, burying behavior is not simply a reflection of a repetitive or compulsive phenotype, and instead should be regarded as a unique class of behaviors.

## Materials and Methods

### Experimental procedures

All experimental animal procedures were approved a priori by an independent animal ethical committee (DEC-Consult, Soest, The Netherlands), as required by Dutch law and conform to the relevant institutional regulations of the Erasmus MC, the Netherlands Institute for Neuroscience KNAW, and Dutch legislation on animal experimentation (CCD approval: AVD1010020197846, AVD101002016791, and AVD801002015126).

### Animals

We used male and female mice of the following strains: (1) *Ube3a^m-/p+^* (*Ube3a^tm2Yelg^*; Wang et al., 2018) and their wildtype littermates; (2) *Shank2*^-/-^ mice and their wildtype littermates (Schmeisser et al., 2012; Won et al., 2012); (3) *Sapap3*^-/-^ mice and their wildtype littermates (Welch et al., 2007). Strain 1 was generated in the F1 hybrid 129S2-C57BL/6J background. Strains 2 and 3 were bred on a C57BL/6J background.

Strains 1-2 were between 8 and 12 weeks of age and were housed and tested in the Erasmus MC, Rotterdam, the Netherlands. Animals had ad libitum access to water and food (standard laboratory chow) and were kept on a regular 12-h light/dark cycle. *Shank2*^-/-^ mice and their wildtype littermates were group-housed (3 mice per cage, mixed genotypes in the same cage). *Ube3a^m-/p+^* mice and their wildtype littermates were group-housed (3 mice per cage, mixed genotypes in the same cage). All mice from strains 1-2 were kept on wood chip bedding (Lignocel® Hygienic Animal Bedding, JRS) that was also used for the experiments.

*Sapap3*^-/-^ mice as well as their wildtype littermates (strain 3) were between 8 and 18 weeks of age and were housed and tested at the Netherlands Institute for Neuroscience, Amsterdam, the Netherlands. Animals had *ad libitum* access to water and food (standard laboratory chow) and were group housed on a regular 12-h light/dark cycle. The *Sapap3*^-/-^ mice and their littermates were kept on corn-cob bedding (Bio Services® EuroCob Corn) that was also used for the experiments.

### Behavioral testing

Animals were habituated to the testing room for at least 1 h before experiments. To isolate external factors, all experiments were done inside a 130 x 80 x 80 cm wooden box with a door. The 6 mm high-pressure laminate walls were lined with acoustic foam to reduce external noise penetration. Testing was done in a 26.6 x 42.5 x 18.5 cm cage (Eurostand 1291H-Type III H) which rested on an elevated 10 mm frosted perspex® shelf. The apparatus was evenly lit from top and bottom using white LED strips and recorded with an overhead camera (Basler acA 1300-600gm) with a 4.4-11mm/F1.6ens (KOWA) at 25 frames per second. Testing cages were filled with wood chip bedding (Lignocel® Hygienic Animal Bedding, JRS) for strains 1-2, and corn-cob bedding (Bio Services® EuroCob Corn) for strain 3, to the height of ~4 cm. Next, 20 blue glass marbles were spaced out evenly in four rows on the bedding. The recordings were started immediately after the animals were placed in the cage, using a custom Bonsai script (Lopes et al., 2015). The animals were left to explore the apparatus for 30 minutes, after which the mice were removed and a top-down image of the marbles was taken. The bedding was discarded and the cages were cleaned with 70% ethanol between each experiment.

### Manual scoring of burying behavior

Manual annotation of four 10-minute videos was done by 4 observers using the open-source software BORIS (Friard & Gamba, 2016), which allowed for frame-by-frame annotation.

### Classifier training

A classifier to study burying behavior was created in the open source, MATLAB (Mathworks, R2018a) based Janelia Automatic Animal Behavior Annotator (JAABA®, v0.6) environment (Kabra et al., 2013). Videos were prepared by cropping raw images to the area of the apparatus using FFMPEG (https://www.ffmpeg.org/). To prepare the data for classifier training, the videos were then tracked using open-source software MATLAB based Mouse Tracker (motr) (Ohayon et al., 2013). All videos were processed and tracked in a batch-mode, significantly decreasing the processing time. The output data from motr was converted to the required format for JAABA by using the function “PrepareJAABA”. Version 0.6.0 of JAABA was obtained from SourceForge (http://jaaba.sourceforge.net/). The classifier was trained on 12343 frames in two videos of *Shank2*^-/-^ and two videos of their wildtype littermates. Not all frames were annotated in order to get a relatively equal distribution of burying and non-burying frames. A minimum bout length of one second was applied for the analysis of burying characteristics. The trained classifier is available upon request. However, in order to obtain maximum accuracy, we recommend training a custom classifier on newly acquired videos as the experimental conditions such as light, cage dimensions, mouse color and bedding can differ across laboratories. The classifier can then be used for all newly acquired videos.

To obtain information regarding movement characteristics, all videos were tracked using open-source software Optimouse (Ben-Shaul, 2017). All videos were tracked in a batch-mode, which significantly decreased the processing time. The (x,y) position data from Optimouse was combined with the frame by frame output of JAABA to create heatmaps of burying topography using a custom-written MATLAB script.

### Image analysis

A custom script to analyze buried marble surface area was created in ImageJ (https://imagej.nih.gov/ij/). The marbles are separated from the background with the use of color thresholding, after which the masked-out surface area can be measured per marble.

### Code accessibility

The code/software described in the paper is freely available online at https://github.com/BaduraLab/Marble-Burying.

### Data processing and statistics

All data was processed using Microsoft Excel and custom MATLAB scripts, on a Windows 10 64-bit computer. Statistical group comparisons were done using Graphpad Prism 8 software. The assumption of normality was tested using the D’Agostino & Pearson test. If the data passed the assumption of normality, a one- or two-tailed t-test was used to compare groups. If the assumption of normality was violated, a one- or two-tailed Mann-Whitney test was used.

## Results

### Classification performance

We collected four videos of marble burying (30 min/25fps each) from two female *Shank2*^-/-^ and two female *Shank2*^-/-^ mice of 14-18 weeks old in our custom-built marble burying setup (see Methods for details). We pre-processed the videos by cropping them to the size of the marble burying arena using FFMPEG (https://www.ffmpeg.org/) and subsequently tracked the mice using open source Mouse Tracker (motr) (Ohayon et al., 2013). Next, we transferred the videos to the Janelia Automatic Animal Behavior Annotator (JAABA®) environment (Kabra et al., 2013) and trained the classifier to discriminate the burying events. The classifier achieved the classification accuracy of 83.1% for correctly annotating burying frames and 83.9% for non-burying frames (**Fig. 1A**). Manual annotation of four 10-minute videos by four independent observers blinded to genotype showed high variability in average bout duration between observers (**Fig. 1B**). There was a significant overlap in the annotation made by the observers and the classifier (**Fig. 1C**). Overall, the observers annotated less frames as burying-positive than the classifier (Fig. **1D**).

**Figure 1.**
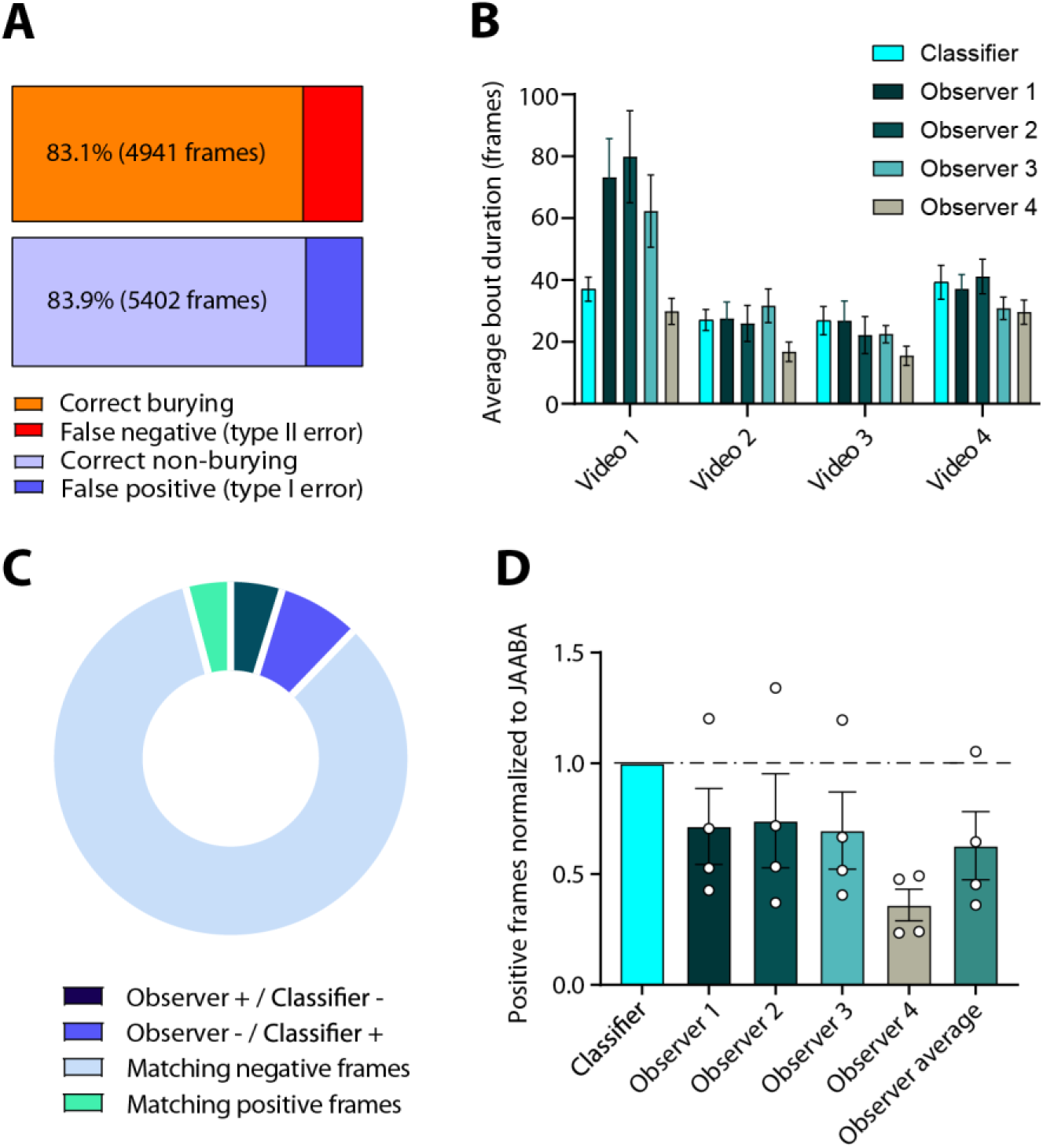
Classifier validation. **A)** Cross-validation results of 12343 manually-annotated frames. The orange bar represents correct annotation of frames containing burying, while red represents frames incorrectly scored as burying by the classifier. Light-blue bar represents correct annotation of non-burying frames, whereas dark-blue shows incorrectly annotated non-burying frames. **B)** Average duration of bouts in four videos of 10 minutes in duration, manually scored by four observers. **C)** Pie chart depicts (1) frames that observers scored as burying whereas the classifier scored non-burying, (2) frames that observers scored as non-burying whereas the classifier scored burying, and (3/4) non-burying and burying frames where observers were in consensus with the classifier. **D)** Frames scored positive for burying by the observers normalized to the results of the classifier. Each dot represents a single video annotated by an observer.

### Classification validation in the Angelman mouse model

In order to validate the results from the classifier, we chose a mouse model with a well-documented phenotype in the marble burying test. *Ube3a* mutant mice (*Ube3a^m-/p+^*) are a model for Angelman Syndrome (AS), which have consistently shown impaired marble-burying behavior (Sonzogni et al., 2018; Rotaru et al., 2020).

All videos were collected and processed as described above and in the Methods. Using our classifier, we indeed found that *Ube3a^m-/p+^* mice (n = 12) spent less time burying than their wildtype littermates (n = 11; p = 0.0175), which was caused by a trend in the decreased number of bouts (p =0.073) as well as shorter average bout duration (p = 0.0405; **Fig. 2A and B**). Next, we used open-source software Optimouse (Ben-Shaul, 2017) to analyze movement characteristics independent of the burying behavior. *Ube3a^m-/p+^* mice travelled significantly less distance (p = 0.0011) during the marble burying test and were significantly slower (n = 12 both genotypes; p = 0.0004; **Fig. 2C**), showing decreased locomotor activity consistent with previous findings (Sonzogni et al., 2018). By combining the output of the tracking software with the output of the classifier we created heatmaps illustrating spatial information regarding burying events. Both *Ube3a^m-/p+^* mice as well as their wildtype littermates had a strong preference for moving in the corners of the apparatus (**Fig 2D**, **top**). However, whilst the WT mice predominantly buried in the corners, the burying behavior of *Ube3a^m-/p+^* mice was less spatially specific (**Fig 2D, bottom**).

**Figure 2.**
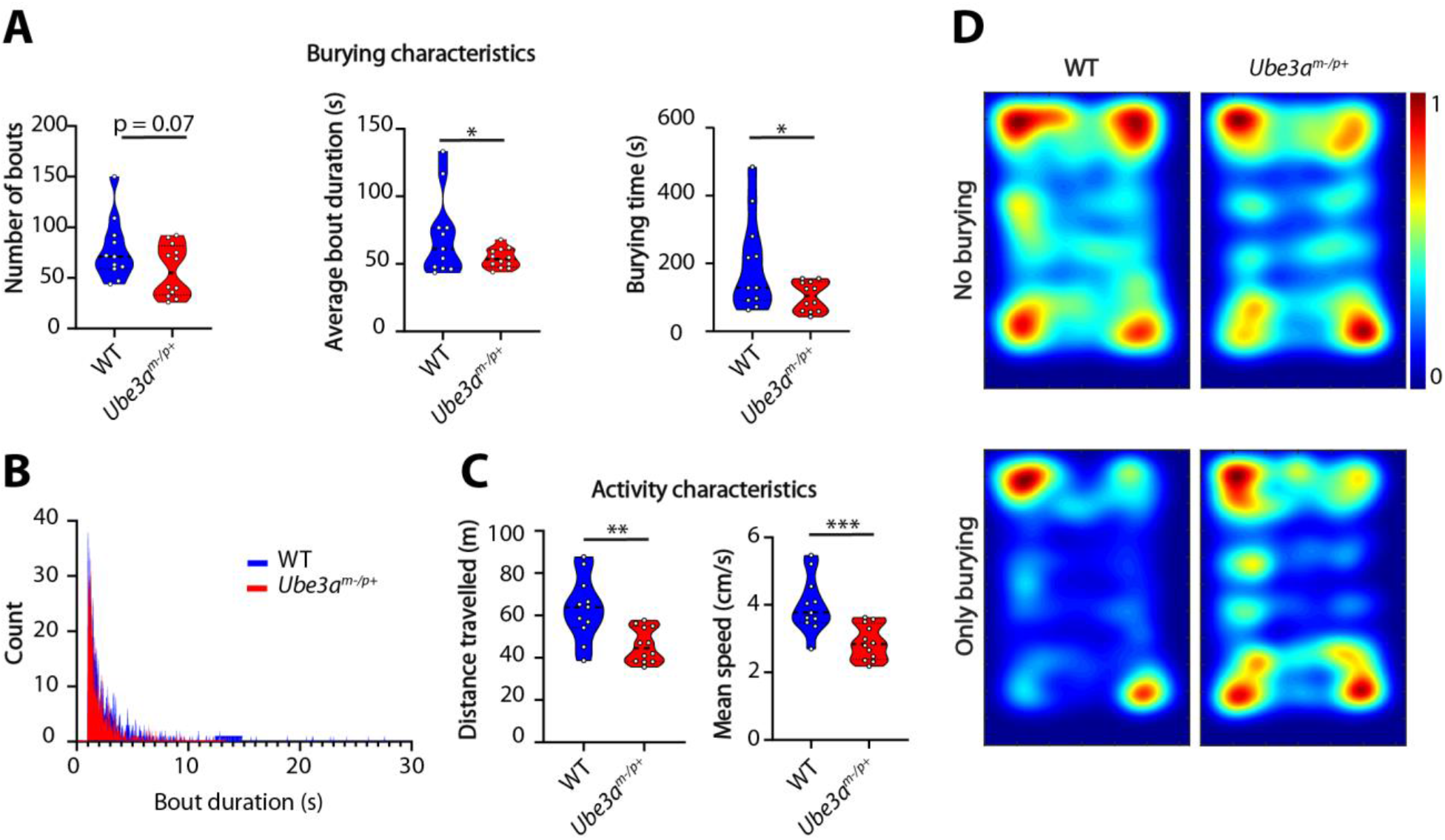
Ube3a mutants show a burying phenotype consistent with literature. **A)** Number of bouts, average bout duration, and total burying time for wildtype and Ube3a^m-/p+^ mice during the marble burying test. Data presented as median with interquartile range (number of bouts, one-tailed Mann-Whitney test; average bout duration and total burying time, two-tailed, unpaired t-test). **B)** Histogram showing distribution of bout lengths and their frequency. **C)** Distance travelled and mean speed over the duration of the test. Data presented as median with interquartile range (one-tailed, unpaired t-test). **D)** Heatmaps showing all frames where mice do not bury (top) and frames where mice show burying behavior (bottom). * p≤0.05, ** p≤0.01, *** p≤0.001, n = 12 for mice per genotype except for the WT group where n = 11 for **A**, **B** and **D** due to erroneous motr tracking coming from an artifact of the experimenter’s hand in the field of view.

### Quantification of complex parameters reveals unique features of marble burying behavior

Burying behavior in the marble burying test is often ascribed to anxiety-like behaviors and/or repetitive behaviors, but conclusive evidence for either type of behavior is lacking. Because our methodological approach enables measuring several behavioral parameters in one assay, we performed the marble burying experiments with two additional mouse models to study this further.

We first tested mice with a mutation in the *Shank2* gene, an established mouse model for ASD (Eltokhi et al., 2018; Kim et al., 2018; Peter et al., 2016) displaying increased repetitive behavior in the grooming assay and hyperactivity (Schmeisser et al., 2012). In the marble burying test however, *Shank2*^-/-^ mice (n = 10 mice per genotype) showed a significant decrease in the number of burying bouts (p = 0.0032) and total burying time (p = 0.0309; **Fig. 3A**). There was no difference in average bout length (p = 0.2894), reflected in no visible difference in shape of the distribution of bout durations (**Fig 3B**). Therefore, we can conclude that the decrease in the number of burying bouts was evenly distributed across shorter and longer bouts. The hyperactivity phenotype was clearly present in the marble burying test as *Shank2* mutants travelled larger distances (p = 0.0005) at a higher speed (p = 0.0008; **Fig. 3C**). *Shank2*^-/-^ mice hyperactivity was evident from the heatmaps, with minor differences in spatial distribution of the burying events when compared to wildtype littermates (**Fig. 3D**).

**Figure 3.**
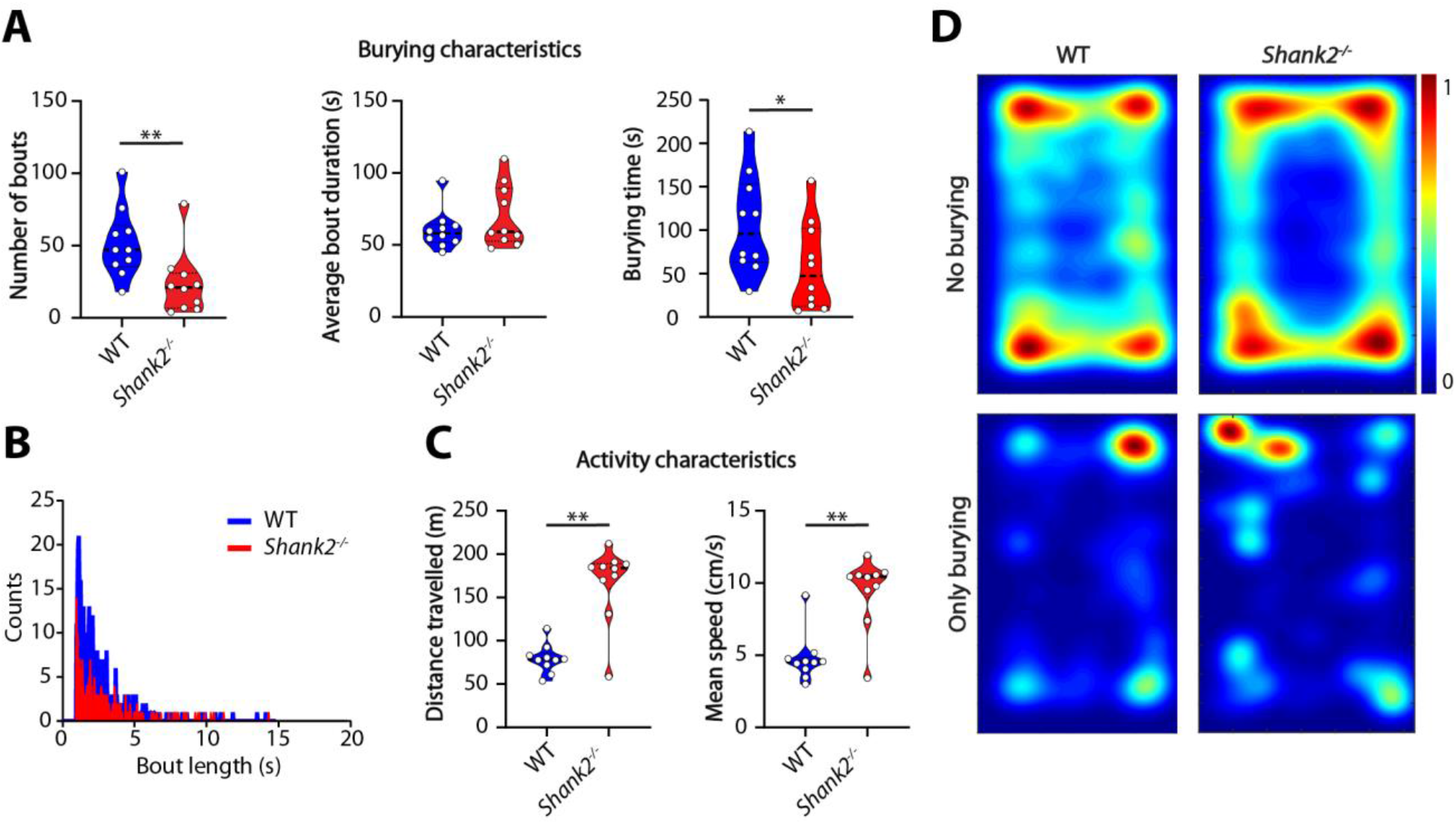
Shank2 mutants show decreased number of bouts but similar bout duration compared to wildtypes. **A)** Number of bouts, average bout duration, and total burying time shown for wildtype and Shank2 mutant mice during the marble burying test. Data presented as median with interquartile range (number of bouts and average bout duration, one-tailed Mann-Whitney test; burying time, one-tailed, unpaired t-test). **B)** Histogram showing distribution of bout lengths and their frequency. **C)** Distance travelled and mean speed over the duration of the test. Data presented as median with interquartile range (one-tailed Mann-Whitney test). **D)** Heatmaps showing all frames where mice do not bury (top) and frames where mice do bury (bottom). * p≤0.05, ** p≤0.01; n = 10 mice per genotype.

We next examined the performance of *Sapap3^-/-^* mice in the marble burying test. *Sapap3^-/-^* mice present with a phenotype that matches considerably with OCD patients, including compulsive grooming, decreased cognitive flexibility, altered habit formation, and increased anxiety-like behavior (Boom et al., 2019; Ehmer, Feenstra, et al., 2020; Welch et al., 2007). However, in the marble burying assay the *Sapap3^-/-^* mice did not show altered burying characteristics: there was no difference between the mutant mice and their wildtype littermates (n = 10 mice per genotype) in number of bouts (p = 0.9441), bout duration (p = 0.9856), total time spent burying (p = 0.9362) or bout-duration distribution (**Fig. 4A and B**). *Sapap3*^-/-^ mice showed a tendency to travel less and at lower speeds (**Fig. 4C**). Consistent with the anxiety phenotype (Welch et al., 2007) they spent most time in the corners (**Fig. 4D, top**), but the spatial distribution of the burying events was not affected.

**Figure 4.**
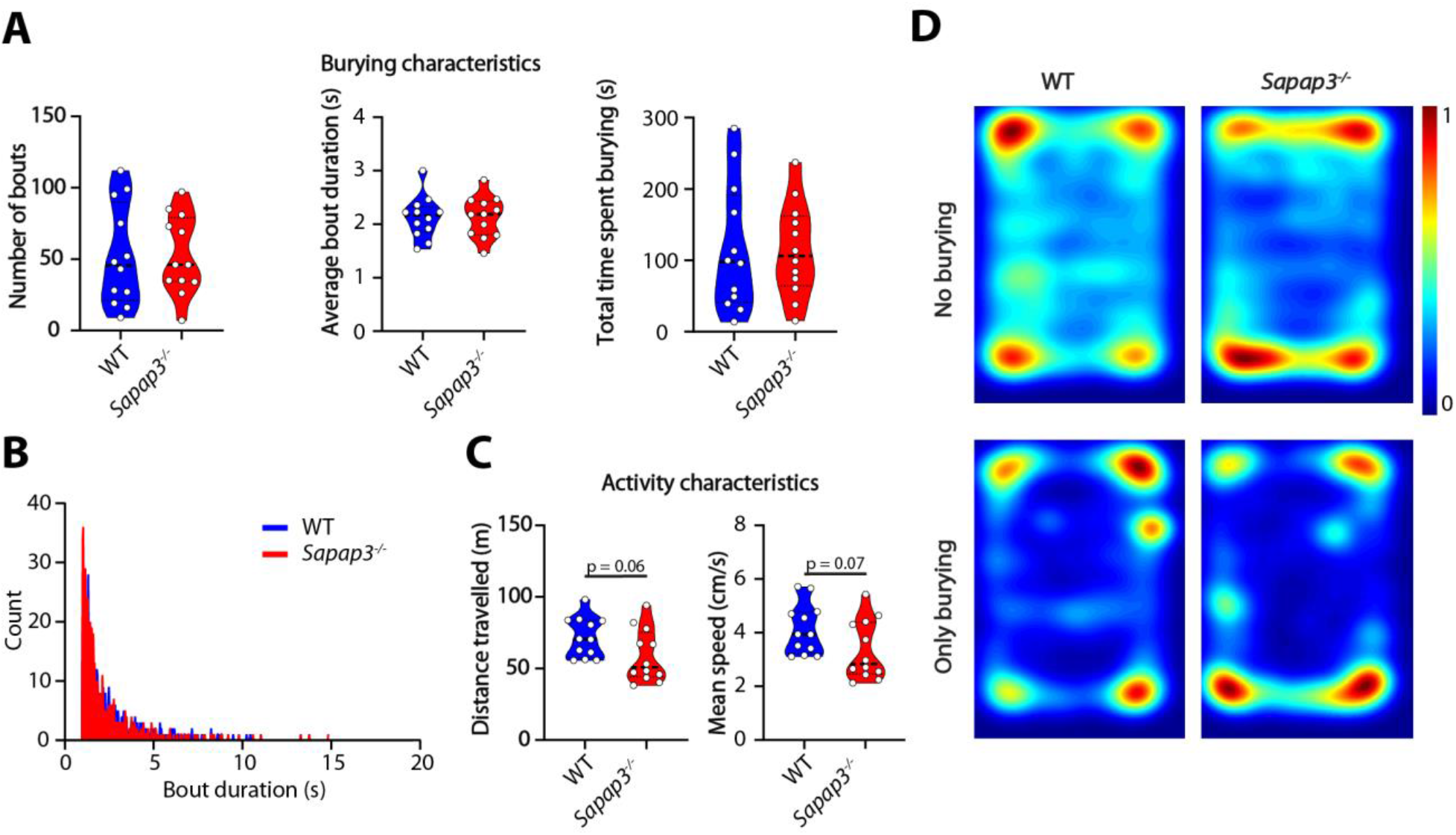
Sapap3 mutants do not show a burying phenotype but tend to travel less distance and do so at decreased speeds. **A)** Number of bouts, average bout duration, and total burying time shown for wildtype mice and Sapap3 mutant mice during the marble burying test. Data presented as median with interquartile range (two-tailed, unpaired t-test). **B)** Histogram showing distribution of bout lengths and their frequency. **C)** Distance travelled and mean speed over the duration of the test. Data presented as median with interquartile range (two-tailed, unpaired t test). **D)** Heatmaps showing all frames where mice do not bury (top) and frames where mice do bury (bottom). n = 10 mice per genotype.

Our methodological approach allows us to quantify the duration and characteristics of the burying events as well as their distribution in time (**Fig 5**). *Ube3a^m-/p+^* and *Shank2*^-/-^ mice showed a consistent decrease in burying behavior over the entire session (**Fig. 5A and B**) compared with control mice. No difference was found for *Sapap3*^-/-^ mice (**Fig. 5C**).

**Figure 5.**
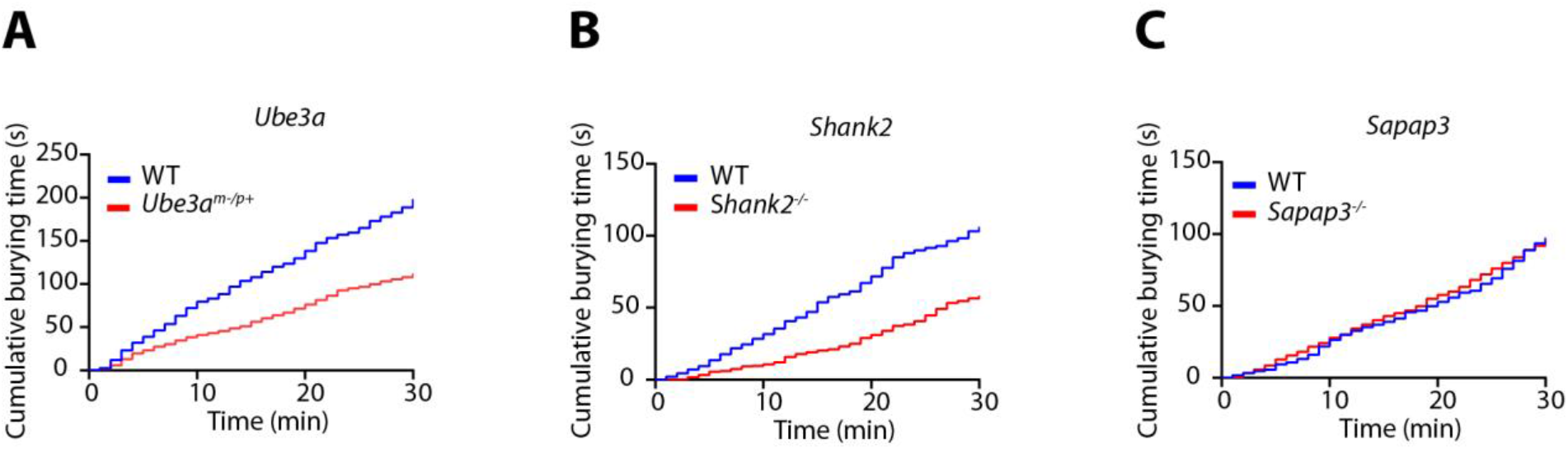
Cumulative burying over time. Time-binned plot with cumulative burying over time. Each bin represents a 1-minute time period. Groups shown are Ube3a^m-/p+^ (**A**), Shank2^-/-^ (**B**) and Sapap3^-/-^ (**C**), and their respective control littermates. n = 12 mice per genotype for **A** and n = 10 per genotype for **B** and **C**.

### Comparison of analysis methods

In order to provide a standardized alternative, fast way of analyzing the result of the marble burying test other than visual scoring, we developed a script in ImageJ to measure buried surface area per marble, based on color thresholding of the marbles and measuring the masked surface area (**Fig. 6A**). We found that the results of the *Ube3a^m-/p+^* mice and their wildtype littermates were comparable between the classifier (p = 0.0175; **Fig. 6B**), experienced visual experimenter scoring (p = 0.0061; **Fig. 6C**), and image analysis (p = 0.0056; **Fig. 6D**). Visual scoring results per animal showed a strong correlation with the analysis of post-burying images using ImageJ (**Fig. 6E, top**). However, burying behavior scored by the classifier showed no direct correlation with the number of buried marbles (**Fig. 6E, bottom**). Although no information is gained over burying characteristics over time, measuring buried marble surface area with ImageJ provides a way to analyze the marble burying result in a way that is similarly fast as visual scoring, yet with replicable results that are not based on experimenter performance or experience.

**Figure 6.**
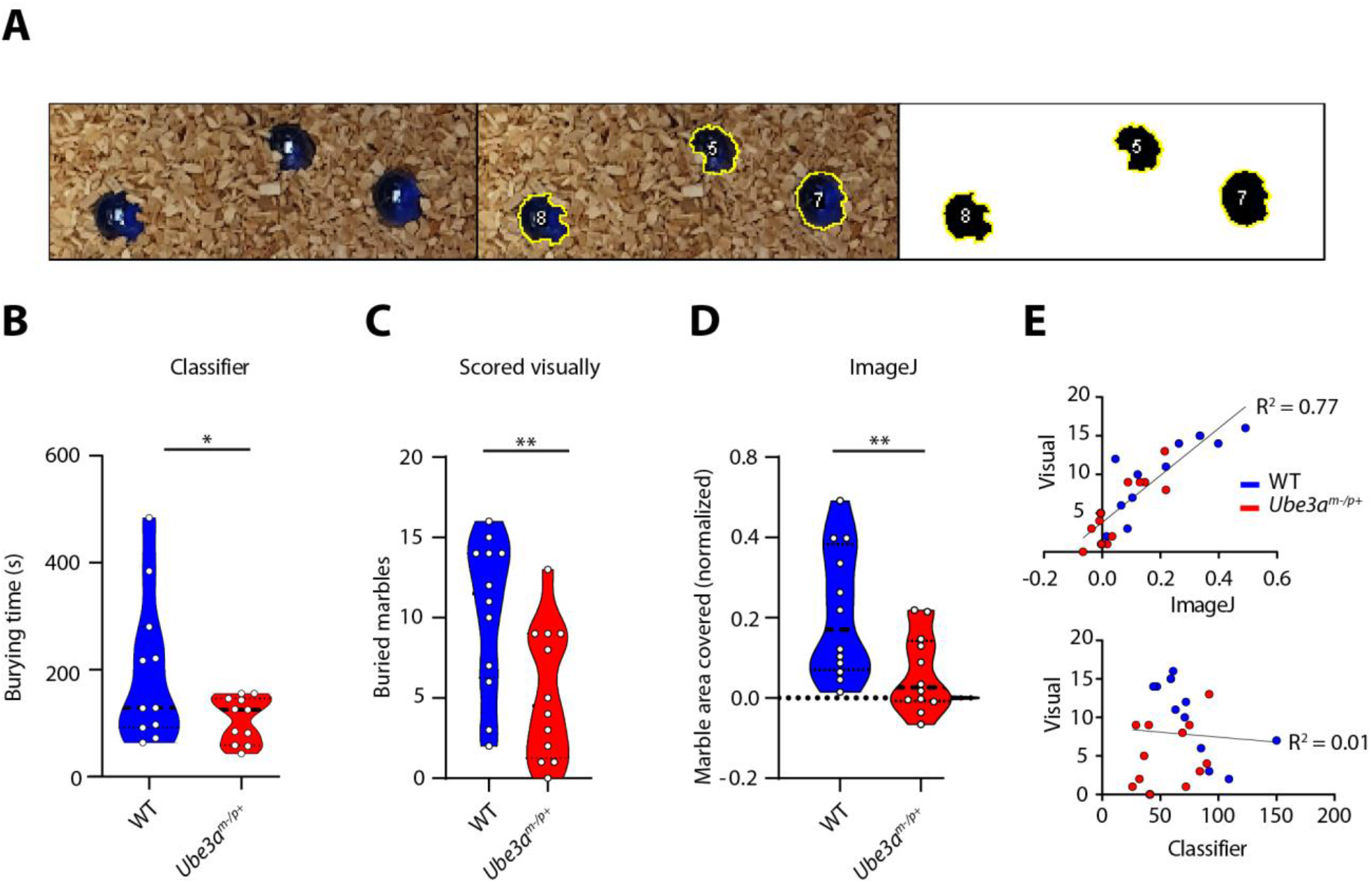
Analysis method comparison. **A)** Example of thresholded marbles analyzed with the ImageJ script. Analysis was done by taking a photo of the buried marbles at the end of each test (left), color thresholding the marbles (middle) and measuring the masked surface area (right). **B)** Ube3a^m-/p+^ results as analyzed with the trained classifier. Data presented as median with interquartile range (one-tailed, unpaired t-test). **C)** Same mice as in A, but analyzed using visual scoring at the end of each test by the experimenters. Data presented as median with interquartile range (one-tailed, unpaired t-test). **D)** Same mice as in A and B, but analyzed with the ImageJ script. Shown is the marble area that is left uncovered. Data presented as median with interquartile range (one-tailed, unpaired t-test). **E)** Scatterplots showing correlation between ImageJ and visual scoring (top) and correlation between the classifier and visual scoring (bottom). * p≤0.05, ** p≤0.01; **A**; n = 11 for WT and n = 12 for Ube3a^m-/p+^; **B** and **C**, n = 12 per genotype).

## Discussion

The marble burying test is often used as an indicator of anxiety and obsessive-compulsive disorders (Angoa-Pérez et al., 2013; Borsini et al., 2002; Broekkamp et al., 1986; Eltokhi et al., 2018; Thomas et al., 2009). However, the meaning of marble burying behavior is highly debated throughout literature. Here, we introduce an analysis method that can increase inter- and intra-experimenter repeatability and establishes marble burying as its own unique behavior. We found that manual annotation of four 10-minute-long videos by four independent observers blinded for genotype showed high variability in the identified average bout duration between observers, indicating that besides being time-consuming, manual annotation of burying bouts lacks reproducibility (**Fig. 1B**). We successfully trained a JAABA classifier, which scored burying behavior in *Ube3a^m-/p+^* mice consistent with existing literature (**Fig. 2A**).

A substantial benefit of automated classification is to allow for non-experienced observers to score the marble burying results in a replicable manner that is consistent with skilled experimenter scoring the number of buried marbles (**Fig. 6**). Our classifier was consistent and generalizable across varying laboratory settings, with differences in the bedding materials and behavioral boxes (wood chip bedding for *Ube3a* and *Shank2* groups, and corn cob bedding for *Sapap3* mice), which demonstrates its applicability across different behavioral setups.

Additional information of mouse behavioral patterns gained from automated classification is an important step towards elucidating the biological meaning of marble burying. While a visual quantification of buried marbles at the end of each test results in a single output parameter (i.e. number of marbles buried), characterizing the actual behavior provides insight into specific burying characteristics and varying burying patterns over time. For example, *Sapap3*^-/-^ mice were previously shown to have a significant shift to longer grooming events and differentiating grooming probability over time (Ramírez-Armenta et al., 2021), however we did not observe a similar pattern in the marble burying test (**Fig. 5C**). In contrast, we found that the *Ube3a^m-/p+^* and *Shank2^-/-^* mice showed a decrease in burying behavior over the entire duration of the test (**Fig. 5A and B**). This analysis is suitable for testing pharmacological, chemogenetic and optogenetic interventions, where tracking changes in the behavior over time is a crucial experimental output. Finally, the use of the classifier in combination with the tracking software reveals novel spatial-information parameters, previously unavailable in the marble burying test. Our analysis showed a striking decrease in *Ube3a^m-/p+^* specificity for burying in the corners of the arena (**Fig. 2D**). Overall, the activity analysis shows that there is no correlation between the level of activity and burying behavior as *Shank2^-/-^* mice show clear hyperactivity but less burying bouts compared to their wildtype littermates. This finding is in contrast to a recently published study (Berg et al., 2021), where the authors discuss low activity as a potential cause for decreased number of buried marbles in an Angelman Syndrome mouse model. Our observations are however consistent with the recent study showing that rescue of the *Ube3a* expression in the juvenile mice alleviates the motor deficits in the rotarod assay but not the marble burying (Milazzo et al., 2021). Together this data strongly indicates that motor activity is not directly related to the burying behavior.

For cases in which additional information about burying characteristics is not required, we introduced a custom-written ImageJ script for the analysis of buried marble surface area (**Fig. 6A**). This method showed consistent results with both the classifier (**Fig. 6B**) and visual scoring of buried marbles (**Fig. 6C**), and showed a strong correlation with visual scoring results (**Fig. 6E, top**). Burying behavior showed no clear correlation with the number of buried marbles on a per-animal basis (**Fig. 6E, bottom**). This disparity can be caused by factors such as marbles being buried and unburied several times over the duration of the test, which cannot be captured by the scoring of the resulting buried marbles. The lack of correlation between the burying events and the number of buried marbles further emphasizes the need for using the methods, such as our classifier, that directly measure the burying behavior instead of focusing solely on the outcome (buried marbles).

By combining the additional information gained from the classifier with selective mouse models known for repetitive and compulsive behaviors we aimed to address the paucity of understanding the implications of the marble burying test. Previous studies have shown increased repetitive behavior and an obsessive-compulsive phenotype in *Shank2*^-/-^ and *Sapap3*^-/-^ mice, represented by the increased levels of self-grooming behavior (Schmeisser et al., 2012; Welch et al., 2007). In our experiments, *Shank2*^-/-^ mice showed a significant decrease in the number of burying bouts and overall burying time (**Fig. 3A**). No difference was found between *Sapap3*^-/-^ mice and their wildtype littermates (**Fig. 4**). These results indicate that compulsivity is not a unitary construct (Ehmer, Crown, et al., 2020) and that burying behavior, although it is often ascribed to repetitive behaviors (Thomas et al., 2009), captures a distinctly different behavioral aspect than self-grooming. Grooming is an innate behavior that is maintained in the absence of sensory feedback and when it is no longer serving its regular maintenance function (Spruijt et al., 1992). However, burying behavior integrates sensory stimuli such as spatial information regarding the marble position, suggesting that marble burying belongs to a more complex class of behaviors. Assigning the marble burying behavior to its unique class, distinct from the grooming behavior, can explain not only our own results but many previously published confounding findings. For example, global *Pten* haploinsufficient mice (an ASD mouse model) show increased levels of repetitive grooming as well as increased levels of burying behavior (Clipperton-Allen & Page, 2014). However, the neuronal-specific *Pten* knock-out mice present with decreased levels of burying (Lugo et al., 2014). Of note, the neuronal-specific *Pten* mutant mice display severe sclerosis of the pyramidal cell layer of the CA3 of the hippocampus (Backman et al., 2001), which further supports the hypothesis that the hippocampus might be important for the spatial aspect of the marble burying test. More research is necessary to elucidate the exact biological background of burying behavior.

## Conclusion

In this study we provide a novel method for replicable analysis of the marble burying test by automated classification of behavior. The classifier scored decreased levels of *Ube3a^m-/p+^* burying behavior consistent with literature, providing a way to reduce inter- and intra-experimenter variability as well as allowing non-experienced observers to accurately analyze the marble burying test. We provide a reproducible alternative in the form of an image analysis script for cases in which additional information is not required. Mouse models of ASD and OCD, which previously showed increased levels of compulsive-like behavior, were found to have decreased levels of burying behavior (*Shank2*^-/-^) or no difference from wildtypes (*Sapap3*^-/-^) which establishes marble burying as a distinct behavior different from repetitive behaviors such as grooming.

## Acknowledgments

We thank Roxanne ter Haar for assistance with breedings and animal experiments. We would also like to thank Bastijn van den Boom for the valuable discussions.

## References

Angoa-Pérez, M., Kane, M. J., Briggs, D. I., Francescutti, D. M., & Kuhn, D. M. (2013). Marble Burying and Nestlet Shredding as Tests of Repetitive, Compulsive-like Behaviors in Mice. Journal of Visualized Experiments: JoVE, 82, 50978. https://doi.org/10.3791/50978

Aso, Y., Sitaraman, D., Ichinose, T., Kaun, K. R., Vogt, K., Belliart-Guérin, G., Plaçais, P.-Y., Robie, A. A., Yamagata, N., Schnaitmann, C., Rowell, W. J., Johnston, R. M., Ngo, T.-T. B., Chen, N., Korff, W., Nitabach, M. N., Heberlein, U., Preat, T., Branson, K. M., … Rubin, G. M. (2014). Mushroom body output neurons encode valence and guide memory-based action selection in Drosophila. ELife, 3, e04580. https://doi.org/10.7554/eLife.04580

Backman, S. A., Stambolic, V., Suzuki, A., Haight, J., Elia, A., Pretorius, J., Tsao, M.-S., Shannon, P., Bolon, B., Ivy, G. O., & Mak, T. W. (2001). Deletion of Pten in mouse brain causes seizures, ataxia and defects in soma size resembling Lhermitte-Duclos disease. Nature Genetics, 29(4), 396–403. https://doi.org/10.1038/ng782

Ben-Shaul, Y. (2017). OptiMouse: A comprehensive open source program for reliable detection and analysis of mouse body and nose positions. BMC Biology, 15(1), 41. https://doi.org/10.1186/s12915-017-0377-3

Berg, E. L., Petkova, S. P., Born, H. A., Adhikari, A., Anderson, A. E., & Silverman, J. L. (2021). Insulin-like growth factor-2 does not improve behavioral deficits in mouse and rat models of Angelman Syndrome. Molecular Autism, 12(1), 59. https://doi.org/10.1186/s13229-021-00467-1

Blut, C., Crespi, A., Mersch, D., Keller, L., Zhao, L., Kollmann, M., Schellscheidt, B., Fülber, C., & Beye, M. (2017). Automated computer-based detection of encounter behaviours in groups of honeybees. Scientific Reports, 7(1), 17663. https://doi.org/10.1038/s41598-017-17863-4

Boom, B. J. G. van den, Mooij, A. H., Misevičiūtė, I., Denys, D., & Willuhn, I. (2019). Behavioral flexibility in a mouse model for obsessive-compulsive disorder: Impaired Pavlovian reversal learning in SAPAP3 mutants. Genes, Brain and Behavior, 18(4), e12557. https://doi.org/10.1111/gbb.12557

Boom, B. J. G. van den, Pavlidi, P., Wolf, C. J. H., Mooij, A. H., & Willuhn, I. (2017). Automated classification of self-grooming in mice using open-source software. Journal of Neuroscience Methods, 289, 48–56. https://doi.org/10.1016/j.jneumeth.2017.05.026

Borsini, F., Podhorna, J., & Marazziti, D. (2002). Do animal models of anxiety predict anxiolytic-like effects of antidepressants? Psychopharmacology, 163(2), 121–141. https://doi.org/10.1007/s00213-002-1155-6

Broekkamp, C. L., Rijk, H. W., Joly-Gelouin, D., & Lloyd, K. L. (1986). Major tranquillizers can be distinguished from minor tranquillizers on the basis of effects on marble burying and swim-induced grooming in mice. European Journal of Pharmacology, 126(3), 223–229. https://doi.org/10.1016/0014-2999(86)90051-8

Çalişkan, H., Şentunali, B., Özden, F. M., Cihan, K. H., Uzunkulaoğlu, M., Çakan, O., Kankal, S., & Zaloğlu, N. (2017). Marble Burying Test Analysis in Terms of Biological and Non-Biological Factors. Journal of Applied Biological Sciences, 11(1), 54–57.

Clipperton-Allen, A. E., & Page, D. T. (2014). Pten haploinsufficient mice show broad brain overgrowth but selective impairments in autism-relevant behavioral tests. Human Molecular Genetics, 23(13), 3490–3505. https://doi.org/10.1093/hmg/ddu057

Ehmer, I., Crown, L., Leeuwen, W. van, Feenstra, M., Willuhn, I., & Denys, D. (2020). Evidence for Distinct Forms of Compulsivity in the SAPAP3 Mutant-Mouse Model for Obsessive-Compulsive Disorder. ENeuro, 7(2). https://doi.org/10.1523/ENEURO.0245-19.2020

Ehmer, I., Feenstra, M., Willuhn, I., & Denys, D. (2020). Instrumental learning in a mouse model for obsessive-compulsive disorder: Impaired habit formation in Sapap3 mutants. Neurobiology of Learning and Memory, 168, 107162. https://doi.org/10.1016/j.nlm.2020.107162

Eltokhi, A., Rappold, G., & Sprengel, R. (2018). Distinct Phenotypes of Shank2 Mouse Models Reflect Neuropsychiatric Spectrum Disorders of Human Patients With SHANK2 Variants. Frontiers in Molecular Neuroscience, 11, 240. https://doi.org/10.3389/fnmol.2018.00240

Friard, O., & Gamba, M. (2016). BORIS: A free, versatile open-source event-logging software for video/audio coding and live observations. Methods in Ecology and Evolution, 7(11), 1325–1330. https://doi.org/10.1111/2041-210X.12584

Huang, H.-S., Burns, A. J., Nonneman, R. J., Baker, L. K., Riddick, N. V., Nikolova, V. D., Riday, T. T., Yashiro, K., Philpot, B. D., & Moy, S. S. (2013). Behavioral deficits in an Angelman syndrome model: Effects of genetic background and age. Behavioural Brain Research, 243, 79–90. https://doi.org/10.1016/j.bbr.2012.12.052

Ichimaru, Y., Egawa, T., & Sawa, A. (1995). 5-HT1A-Receptor Subtype Mediates the Effect of Fluvoxamine, a Selective Serotonin Reuptake Inhibitor, on Marble-Burying Behavior in Mice. Japanese Journal of Pharmacology, 68(1), 65–70. https://doi.org/10.1254/jjp.68.65

Kabra, M., Robie, A. A., Rivera-Alba, M., Branson, S., & Branson, K. (2013). JAABA: Interactive machine learning for automatic annotation of animal behavior. Nature Methods, 10(1), 64–67. https://doi.org/10.1038/nmeth.2281

Kalariya, M., Prajapati, R., Parmar, S. K., & Sheth, N. (2015). Effect of hydroalcoholic extract of leaves of Colocasia esculenta on marble-burying behavior in mice: Implications for obsessive–compulsive disorder. Pharmaceutical Biology, 53(8), 1239–1242. https://doi.org/10.3109/13880209.2015.1014923

Kim, R., Kim, J., Chung, C., Ha, S., Lee, S., Lee, E., Yoo, Y.-E., Kim, W., Shin, W., & Kim, E. (2018). Cell-Type-Specific Shank2 Deletion in Mice Leads to Differential Synaptic and Behavioral Phenotypes. Journal of Neuroscience, 38(17), 4076–4092. https://doi.org/10.1523/JNEUROSCI.2684-17.2018

Lugo, J. N., Smith, G. D., Arbuckle, E. P., White, J., Holley, A. J., Floruta, C. M., Ahmed, N., Gomez, M. C., & Okonkwo, O. (2014). Deletion of PTEN produces autism-like behavioral deficits and alterations in synaptic proteins. Frontiers in Molecular Neuroscience, 7, 27. https://doi.org/10.3389/fnmol.2014.00027

Milazzo, C., Mientjes, E. J., Wallaard, I., Rasmussen, S. V., Erichsen, K. D., Kakunuri, T., van der Sman, A. S. E., Kremer, T., Miller, M. T., Hoener, M. C., & Elgersma, Y. (2021). Antisense oligonucleotide treatment rescues UBE3A expression and multiple phenotypes of an Angelman syndrome mouse model. JCI Insight, 6(15), e145991. https://doi.org/10.1172/jci.insight.145991

Nicolas, L. B., Kolb, Y., & Prinssen, E. P. M. (2006). A combined marble burying–locomotor activity test in mice: A practical screening test with sensitivity to different classes of anxiolytics and antidepressants. European Journal of Pharmacology, 547(1), 106–115. https://doi.org/10.1016/j.ejphar.2006.07.015

Ohayon, S., Avni, O., Taylor, A. L., Perona, P., & Roian Egnor, S. E. (2013). Automated multi-day tracking of marked mice for the analysis of social behaviour. Journal of Neuroscience Methods, 219(1), 10–19. https://doi.org/10.1016/j.jneumeth.2013.05.013

Pereira, T. D., Aldarondo, D. E., Willmore, L., Kislin, M., Wang, S. S.-H., Murthy, M., & Shaevitz, J. W. (2019). Fast animal pose estimation using deep neural networks. Nature Methods, 16(1), 117–125. https://doi.org/10.1038/s41592-018-0234-5

Peter, S., ten Brinke, M. M., Stedehouder, J., Reinelt, C. M., Wu, B., Zhou, H., Zhou, K., Boele, H.-J., Kushner, S. A., Lee, M. G., Schmeisser, M. J., Boeckers, T. M., Schonewille, M., Hoebeek, F. E., & De Zeeuw, C. I. (2016). Dysfunctional cerebellar Purkinje cells contribute to autism-like behaviour in Shank2-deficient mice. Nature Communications, 7(1), 12627. https://doi.org/10.1038/ncomms12627

Pinel, J. P., & Treit, D. (1978). Burying as a defensive response in rats. Journal of Comparative and Physiological Psychology, 92(4), 708–712. https://doi.org/10.1037/h0077494

Ramírez-Armenta, K. I., Alatriste-León, H., Verma-Rodríguez, A. K., Llanos-Moreno, A., Ramírez-Jarquín, J. O., & Tecuapetla, F. (2021). Optogenetic inhibition of indirect pathway neurons in the dorsomedial striatum reduces excessive grooming in Sapap3-knockout mice. Neuropsychopharmacology, 1–11. https://doi.org/10.1038/s41386-021-01161-9

Rotaru, D. C., Mientjes, E. J., & Elgersma, Y. (2020). Angelman Syndrome: From Mouse Models to Therapy. Neuroscience, 445, 172–189. https://doi.org/10.1016/j.neuroscience.2020.02.017

Schmeisser, M. J., Ey, E., Wegener, S., Bockmann, J., Stempel, A. V., Kuebler, A., Janssen, A.-L., Udvardi, P. T., Shiban, E., Spilker, C., Balschun, D., Skryabin, B. V., Dieck, S. tom, Smalla, K.-H., Montag, D., Leblond, C. S., Faure, P., Torquet, N., Le Sourd, A.-M., … Boeckers, T. M. (2012). Autistic-like behaviours and hyperactivity in mice lacking ProSAP1/Shank2. Nature, 486(7402), 256–260. https://doi.org/10.1038/nature11015

Serra, I., Manusama, O. R., Kaiser, F. M. P., Floriano, I. I., Wahl, L., Zalm, C. van der, IJspeert, H., Hagen, P. M. van, Beveren, N. J. M. van, Arend, S. M., Okkenhaug, K., Pel, J. J. M., Dalm, V. A. S. H., & Badura, A. (2021). Activated PI3Kδ syndrome, an immunodeficiency disorder, leads to sensorimotor deficits recapitulated in a murine model (p. 2021.01.15.426862). https://doi.org/10.1101/2021.01.15.426862

Sonzogni, M., Hakonen, J., Bernabé Kleijn, M., Silva-Santos, S., Judson, M. C., Philpot, B. D., van Woerden, G. M., & Elgersma, Y. (2019). Delayed loss of UBE3A reduces the expression of Angelman syndrome-associated phenotypes. Molecular Autism, 10(1), 23. https://doi.org/10.1186/s13229-019-0277-1

Sonzogni, M., Wallaard, I., Santos, S. S., Kingma, J., du Mee, D., van Woerden, G. M., & Elgersma, Y. (2018). A behavioral test battery for mouse models of Angelman syndrome: A powerful tool for testing drugs and novel Ube3a mutants. Molecular Autism, 9, 47. https://doi.org/10.1186/s13229-018-0231-7

Spruijt, B. M., van Hooff, J. A., & Gispen, W. H. (1992). Ethology and neurobiology of grooming behavior. Physiological Reviews, 72(3), 825–852. https://doi.org/10.1152/physrev.1992.72.3.825

Thomas, A., Burant, A., Bui, N., Graham, D., Yuva-Paylor, L. A., & Paylor, R. (2009). Marble burying reflects a repetitive and perseverative behavior more than novelty-induced anxiety. Psychopharmacology, 204(2), 361–373. https://doi.org/10.1007/s00213-009-1466-y

Wang, T., van Woerden, G. M., Elgersma, Y., & Borst, J. G. G. (2018). Enhanced Transmission at the Calyx of Held Synapse in a Mouse Model for Angelman Syndrome. Frontiers in Cellular Neuroscience, 11, 418. https://doi.org/10.3389/fncel.2017.00418

Welch, J. M., Lu, J., Rodriguiz, R. M., Trotta, N. C., Peca, J., Ding, J.-D., Feliciano, C., Chen, M., Adams, J. P., Luo, J., Dudek, S. M., Weinberg, R. J., Calakos, N., Wetsel, W. C., & Feng, G. (2007). Cortico-striatal synaptic defects and OCD-like behaviours in Sapap3-mutant mice. Nature, 448(7156), 894–900. https://doi.org/10.1038/nature06104

Wiltschko, A. B., Tsukahara, T., Zeine, A., Anyoha, R., Gillis, W. F., Markowitz, J. E., Peterson, R. E., Katon, J., Johnson, M. J., & Datta, S. R. (2020). Revealing the structure of pharmacobehavioral space through motion sequencing. Nature Neuroscience, 23(11), 1433–1443. https://doi.org/10.1038/s41593-020-00706-3

Won, H., Lee, H.-R., Gee, H. Y., Mah, W., Kim, J.-I., Lee, J., Ha, S., Chung, C., Jung, E. S., Cho, Y. S., Park, S.-G., Lee, J.-S., Lee, K., Kim, D., Bae, Y. C., Kaang, B.-K., Lee, M. G., & Kim, E. (2012). Autistic-like social behaviour in Shank2-mutant mice improved by restoring NMDA receptor function. Nature, 486(7402), 261–265. https://doi.org/10.1038/nature11208

